# *In vitro* modelling of oral microbial invasion in the human colon

**DOI:** 10.1101/2022.10.17.512642

**Authors:** Lucie Etienne-Mesmin, Victoria Meslier, Ophélie Uriot, Elora Fournier, Charlotte Deschamps, Sylvain Denis, Aymeric David, Sarah Jegou, Christian Morabito, Benoit Quinquis, Florence Thirion, Florian Plaza Oñate, Emmanuelle Le Chatelier, S. Dusko Ehrlich, Stéphanie Blanquet-Diot, Mathieu Almeida

## Abstract

Recent advances in the human microbiome characterization have revealed significant oral microbial detection in stools of dysbiotic patients. However, little is known about the potential interactions of these invasive oral microorganisms with commensal intestinal microbiota and host. In this proof of concept study, we propose a new model of oral to gut invasion by the combined use of an *in vitro* model simulating both the physicochemical and microbial (lumen and mucus-associated microbes) parameters of the human colon (M-ARCOL), a salivary enrichment protocol and whole metagenome shotgun sequencing. Oral invasion of the intestinal microbiota was simulated by injection of enriched saliva in the *in vitro* colon model inoculated with faecal sample from the same healthy adult donor. The mucosal compartment of M-ARCOL was able to retain the highest species richness levels over time, whilst it decreased in the luminal compartment. This study also showed that oral microorganisms preferably colonized the mucosal microenvironment, suggesting potential oral-to-intestinal mucosal competitions. This new model of oral-to-gut invasion can provide useful mechanistic insights into the role of oral microbiome in various disease processes.

## INTRODUCTION

The human gastro-intestinal (GI) tract harbors a vast and complex community including between 10 to 100 trillion of microbes dominated by bacteria, collectively referred to as gut microbiota (1). The gut microbiota is playing a major role in host physiology, with an involvement in energy extraction from food, vitamin synthesis, maturation of the immune system and protection against invasion by enteric pathogens (2, 3). In the human body, oral and GI microbiomes represent the two largest microbial ecosystems (4, 5) and pioneering data from the Human Microbiome Project demonstrated they are taxonomically diverse, representing 26% and 29% of total bacteria from the human body, respectively (6).

Studies have shown that saliva contains approximatively 10^7^ to 10^9^ bacteria per milliter (7, 8) with a global diversity of approximately 700 species listed from the oral cavity of healthy subjects. These species are members of Firmicutes, Proteobacteria, Bacteroidetes, Actinobacteria, Fusobacteria, Spirochaetes, Synergistetesand TM7 (5, 9). *Streptococcus* is the most abundant genus in the oral site, with *Haemophilus, Veillonella* and *Prevotella* also prevalent (5, 10, 11). Countless studies have demonstrated that microbiota composition distinctively changes all along the GI tract due to differences in term of oxygenation, substrates availability, pH and residence time between digestive compartments, therefore inducing microbial species niche preferences (12). The highest bacterial load (10^11^ to 10^12^ bacteria per g) and diversity are reached in the colon where predominant phyla are Bacteroidetes, Firmicutes, Actinobacteria and to a lesser extend Proteobacteria and Verrucomicrobia (13–16). It is now well established that each individual harbors a unique gut microbiota composed of an estimated 300 bacterial species detected per healthy individual on average (17–19).

Despite physical distance and chemical hurdles that keep apart the oral microbiome from the gut microbiome, cumulative evidence supports that the oral microbiota is present in the overall gut microbial ecosystem. Li *et al*. demonstrated that transplantation of human saliva to gnotobiotic mice led to a distribution of oral genera throughout the GI tract, with *Fusobacterium, Haemophilus, Streptococcus* and *Veillonella* being especially abundant in the gut of recipient mice (20). In humans, independent studies have demonstrated that some bacterial genera detected in the same healthy subject can overlap between oral and stool samples, confirming an extensive transmission of microbes through the GI tract (11, 21). Such phenomenon of oral-gut transmission occurring under physiological situation seems to be amplified in pathological context. Orally-derived microorganisms are particularly enriched in patients with altered gut microbiota (perturbation termed dysbiosis) and barrier disruption. In particular, Hu and colleagues showed that oral bacteria are enriched in the faecal microbiota of Crohn disease patients (22), suggesting that the oral cavity might act as a reservoir of opportunistic pathogens with the ability to colonize the gut (23), even more importantly in such susceptible host. Likewise, a large fraction of species enriched in the faecal microbiota of patients with liver cirrhosis or after bariatric surgery are of oral origin (24, 25).

To date, the inter-connections between oral and gut microbiota have not been fully elucidated and mechanisms associated to the gut colonization by oral bacteria are not clear. This can be explained by i) the technical difficulties met when analysing oral microbial samples with high resolution shotgun metagenomic sequencing, due to the high proportions of retrieved host DNA and ii) the lack of relevant models. Clinical trials remain as the gold standard approaches, but are hampered by heavy regulatory, technical and costly constraints. For evident ethical reasons, human gut microbiota studies are usually performed with faecal samples making result interpretation difficult since a direct access to the different segments of the GI tract -especially the colon-is limited. Animal models integrate the host-microbe interactions but translation to the human situation remains limited due to major differences of digestive physiology, oral and gut microbiota between most animal models and human.

A relevant alternative in preclinical studies involves the use of *in vitro* model simulating the human digestive environment. *In vitro* models permit to accurately reproduce the complexity and diversity of the *in vivo* microbial ecosystem (26–28) and were recently optimized to incorporate mucin beads leading to mucosal-configuration of the models, *i.e* Mucosal-Simulator of the Human Intestinal Microbial Ecosystem (M-SHIME) and Mucosal M-Artificial Colon (M-ARCOL) (13, 28–32). In the present study, we investigated oral-to-gut microbial invasion by using the M-ARCOL. Our main goal was to validate our experimental approach using shotgun metagenomic analysis of stools and saliva samples from healthy donors and to assess oral microbial invasion on the luminal and mucosal microenvironments in the simulated human colon.

## RESULTS

### Development of a novel experimental design for oral-to-gut invasion in M-ARCOL bioreactors

In this study, we developed an oral-to-gut invasion model using a one-stage fermentation system (M-ARCOL) set-up to reproduce the main physicochemical parameters (pH, temperature, transit time, nutrient availability) found in the human colon (**Fig. 1)**. M-ARCOL bioreactors, composed of two compartments used to mimic the luminal and mucosal microenvironments, were inoculated with faecal samples from two healthy adults. This study was conducted on fecal samples collected from two healthy donors, chosen to represent “extreme” conditions based on their sex, age and nutritional habits, i.e., donor S1 is a methane producer female (aged 22 years old, flexitarian based diet) while donor S2 is a non-methanogen male (aged 52 years old, western like diet). Gas analysis confirmed that anaerobic conditions were efficiently maintained in the bioreactors, by the sole metabolic activity of gut microbiota without gas flushing, which constitutes a main feature of M-ARCOL model compared to other colonic *in vitro* models (**Fig. S1A, B**). Notably, methane production was detected for one of the donors, indicating presence of methanogenic microorganisms. The three-main short chain fatty acids -SCFAs- (acetate, butyrate and propionate) were also measured in the luminal compartment with ratio similar to that found *in vivo* in the human colon, as previously validated (28) (**Fig. S1B, C**). To allow a combined oral invasion experiment and shotgun sequencing on human saliva samples, we generated enriched microbial saliva samples and inoculated them at days D9 and D10 in the luminal phase of M-ARCOL to evaluate oral-to-gut microbial invasion after gut microbial stabilization in the bioreactors (**Fig. 1**). DNA was extracted from all collected samples, sequenced by shotgun metagenomic. All samples from faecal origins (raw stool, luminal and mucosal samples from M-ARCOL) displayed high mapping rates onto the IGC2 gut gene catalogue (median rates >80%), while not onto the oral catalogue (median near 5%). Salivary samples displayed high mapping rates onto the oral microbial gene catalogue (median >79%) and less than 40% mapping rates onto the gut microbial gene catalogue (**Fig. S2** and **Table S1**). Profiles of atmospheric gases and luminal SCFAs were not modified by addition of enriched saliva (**Fig. S1**).

**FIG 1:**
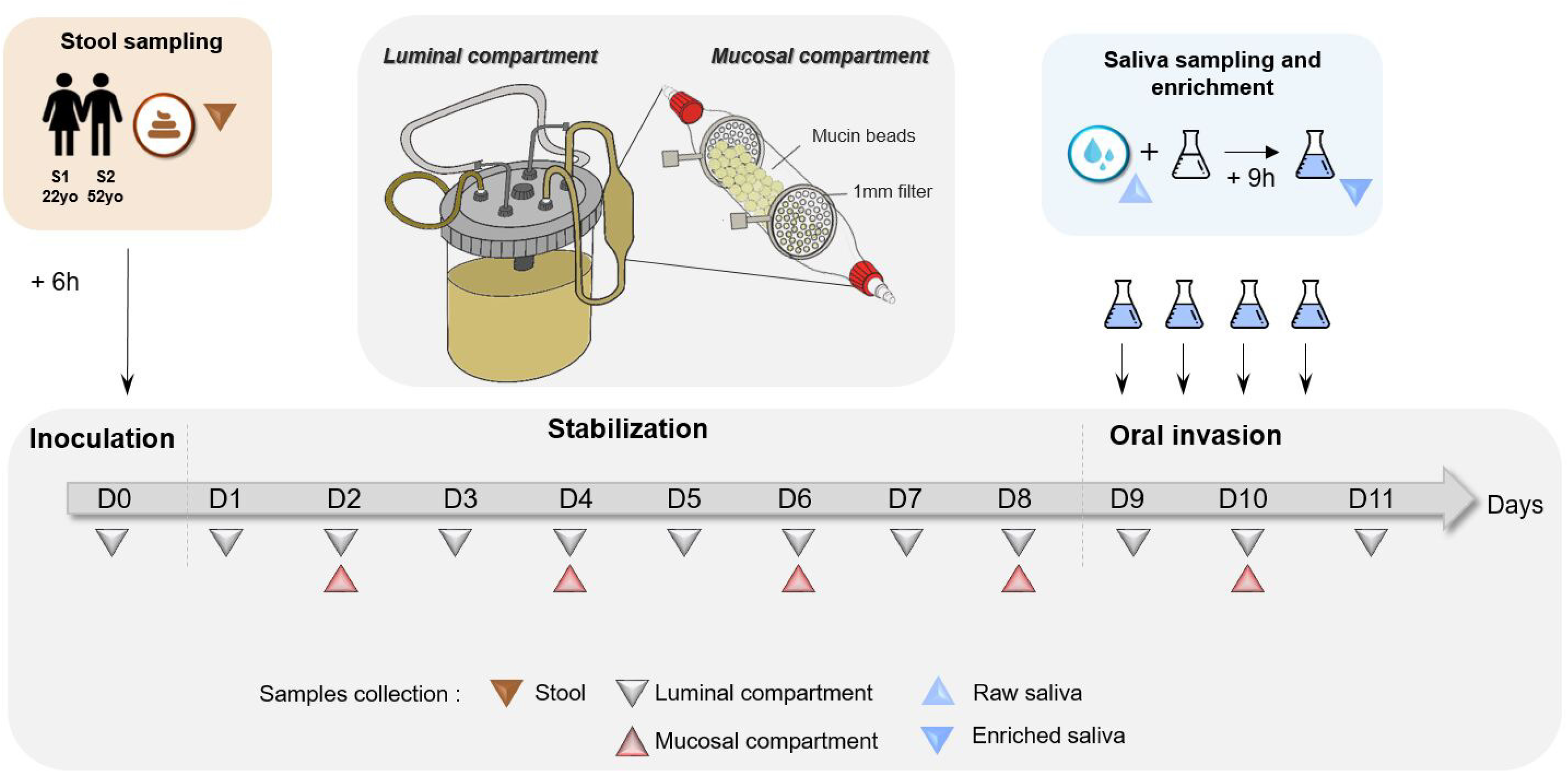
Experimental workflow for *in vitro* fermentation set-up and oral invasion. Fresh stool samples from two healthy donors (S1 and S2) were used to inoculate two independent M-ARCOL bioreactors. Each fermentation was conducted for a total period of 11 days including 24 hours of batch amplification period and 10 days of continuous fermentation. After 8-days stabilization period, oral-to-gut invasion was simulated by injecting a 9-hours enriched saliva from the same donor, twice a day and for two consecutive days (morning and late afternoon of days D9 and D10). Samples were collected from fresh stools (brown triangle), luminal compartment of the bioreactor every day (grey triangle), mucosal compartment every 2 days (red triangle), raw saliva and enriched saliva samples (blue triangle) for each donor.

### The mucosal microenvironment retained higher microbial richness during *in vitro* fermentation

We determined the Metagenomic Species Pangenome (or MSP) richness, defined as clusters of co-abundant genes and representative of microbial species, over time and in the different compartments of the M-ARCOL (**Fig. 2**). For both donors, initial stool and raw saliva samples displayed the highest MSP species richness compared to the bioreactor samples. During *in vitro* fermentations, we observed a loss of richness in the luminal and mucosal compartments, until a stabilization at day D5 in the luminal compartment for both donors. Consistently with faecal MSP richness, S1 donor displayed a higher MSP richness during the first days of fermentation in the luminal and mucosal compartments when compared to S2 donor (delta of 87 and 15 MSP between S1 and S2 donors for stool and raw saliva, respectively). After MSP species richness equilibrium and until the end of the experiment, MSP richness levels were almost equivalent between the two donors in the luminal compartment (delta of 18 MSP at day D11), and individual microbial signature was maintained, as estimated from Bray-Curtis distance measures (**Fig. S3 C**). In the mucosal compartment, MSP richness was found systematically higher than the luminal one at each time point for both donors. The richness loss observed was thus lesser and slower all along the process. The MSP stool richness difference between the donors persisted longer (until at least D8) in the mucosal compartment than in the luminal one (delta of 12 MSP in the luminal and 61 MSP in the mucosal compartment for S1 and S2 donors at day D8). The MSP species richness was found equivalent between S1 and S2 donors in raw saliva samples (delta of 15 MSP), and it decreased in enriched saliva of both donors at days D9 and D10 (delta of 4 MSP between S1 and S2 donors at day D10).

**FIG 2:**
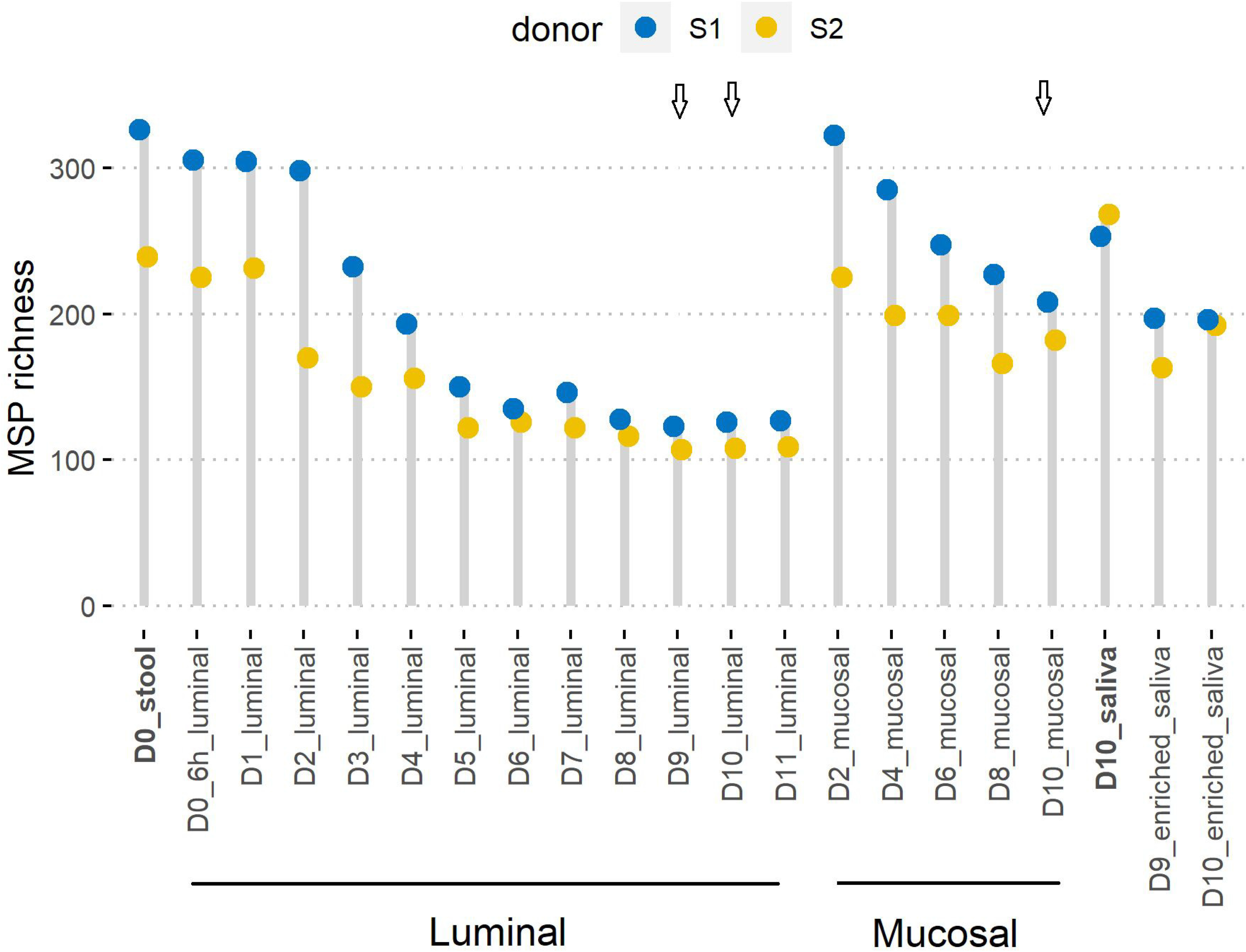
Dot chart for MSP richness dynamic over time. MSP (Metagenomic Species Pangenome) species richness was calculated as the number of detected MSP species in the corresponding sample for donor S1 (blue) and S2 (orange) on the merged MSP abundance table, for luminal and mucosal compartments of the M-ARCOL and for raw saliva and enriched-saliva samples. In bold are indicated initial raw stool and saliva samples. Times of fermentation in the colonic M-ARCOL model are indicated in days (D). Arrows indicate saliva injection into the bioreactors on D9 and D10 days.

### Differential microbial composition between stool, salivary, luminal and mucosal samples

Based on Bray Curtis dissimilarity distance, the overall composition of stool, luminal and mucosal appeared quite close for a given individual and contrasted between the two donors, which is also observed in the salivary microbiota (**Fig. S3**). Indeed, we found that samples clustered together by donor and sample type but also by time points. After the injection of saliva into the bioreactors on days D9 and D10, the luminal and mucosal samples collected were not distant from the same donor faecal samples, suggesting that dominant microbial compositions were not drastically modified by the oral microbial administration. We also observed that raw saliva and enriched saliva clustered together per donor, confirming their close microbial compositions.

We analysed the composition of both saliva, luminal and mucosal associated microbiota in the M-ARCOL at the phylum (**Fig. 3**) and family ranks (**Fig. S4**). At the phylum rank, we found a dominance of Firmicutes and Bacteroidetes followed by Proteobacteria, with some differences between the two donors or between compartments. Main families included *Bacteroidaceae, Rikenellaceae, Prevotellaceae* (Bacteroidetes) across samples and *Ruminococcaceae* or *Streptococcaceae* (Firmicutes) for faecal and oral samples, respectively. Stool of S1 donor was dominated by Firmicutes (75% of relative abundance), stool of S2 donor contained equivalent levels of Bacteroidetes and Firmicutes (0.5 and 0.47 relative abundance, respectively). While the ratio between Firmicutes and Bacteroidetes was modified in the luminal compartment over time, the primary ratio for these taxa observed in stool samples was globally maintained in mucosal samples, particularly for S2 donor. Additional phylum comprised the Verrucomicrobiota phylum (including *Akkermansia muciniphila*), detected in the two donors in the luminal compartment but with higher levels in S1 donor (100 times more – maximum 0.04 relative abundance). The Euryarchaeota phylum (*Methanobrevibacter smithii* species) exclusively detected in S1 donor, coherently with methane detection in gas analyses (**Fig. S1A**), was found in low relative abundance in stool and luminal compartment (<0.01 relative abundance) but was identified in higher levels in the mucosal samples already at day D2 (relative abundance comprised between 0.01 and 0.06). Members of the Actinobacteriodes phylum (mainly of the *Micrococcaceae, Bifidobacteriaceae* and *Actinomycetaceae* families) were found in higher relative abundances in the mucosal and salivary samples (up to 0.02), low in stools samples (below 0.01), and decreased for S1 donor after the oral-to-intestinal invasion. In salivary samples, Firmicutes to Bacteroidetes ratios were found close to those observed in faecal samples for both donors, with additional phyla detected in these samples, such as higher levels of Proteobacteria, Fusobacteria, Patescibacteria and Actinobacteriodes. No major changes in raw and enriched saliva samples were found within each donor, as confirmed by Bray-Curtis dissimilarity analysis (**Fig. S3**).

**FIG 3:**
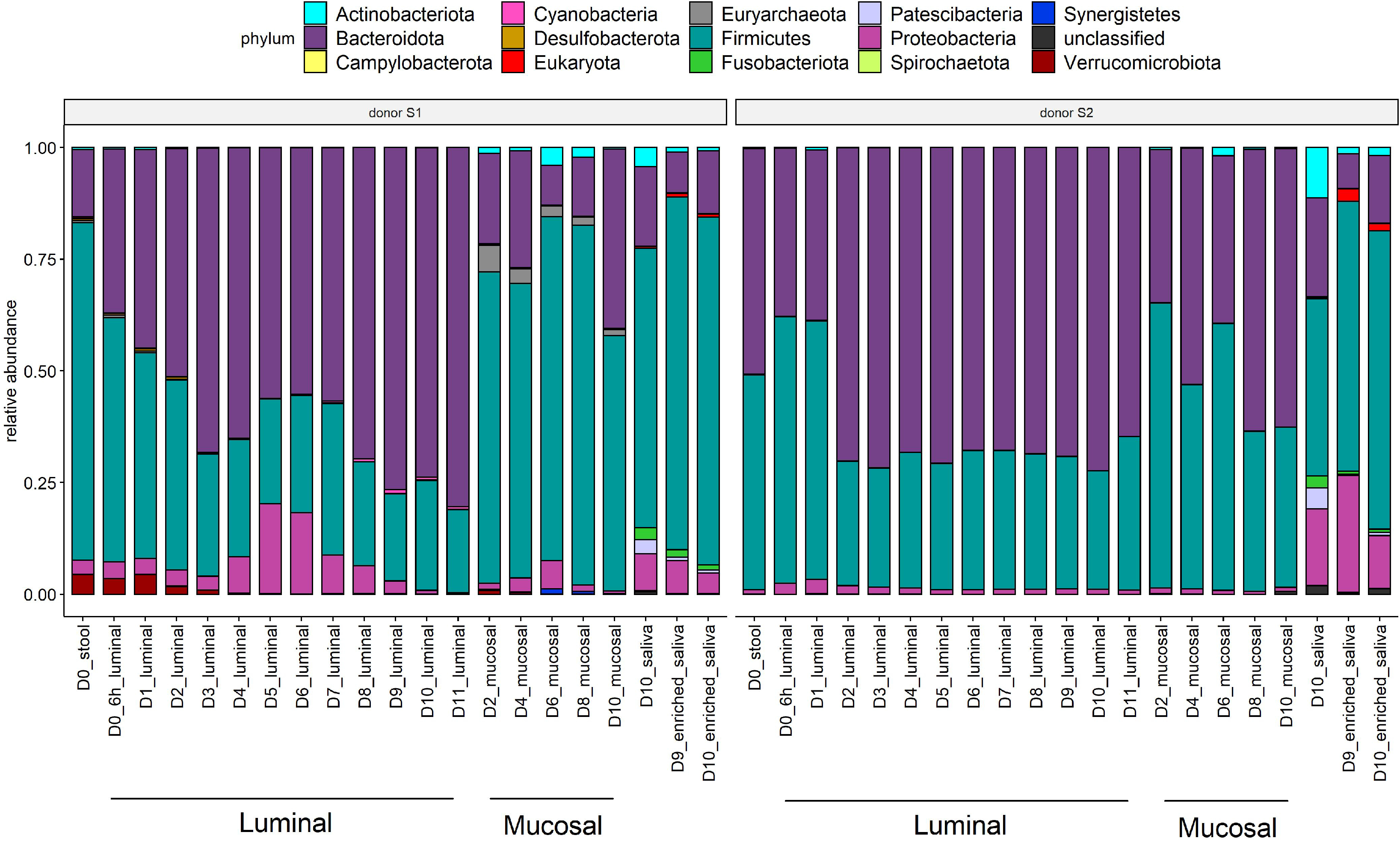
Phylum rank normalized composition per donor. MSP species abundance was normalized per sample by dividing its abundance by the sum of the MSP species abundances detected in the sample. Phylum rank composition was calculated as the sum of the normalized abundances of the corresponding MSP species. Donors (S1 or S2), M-ARCOL compartments (luminal or mucosal) and the days of fermentation are indicated in days (D) are reported.

### A few oral microbial species are present in luminal and mucosal compartments of M-ARCOL before oral invasion

To evaluate the impact of the oral-to-gut invasion on the M-ARCOL microbial composition, we assigned the species to their preferred ecological niche, using their occurrence in raw stools or saliva samples and three species types were defined as: gut, oral or not-determined (ND, because detected in stool and saliva samples, **Table S2**). We then analysed the niche distribution, to assess the number of oral species before and after the invasion in the luminal and mucosal compartments (**Fig. 4**) and the relative abundance of each ecological niche (**Fig. S5**). Raw stools were exclusively dominated by gut-oriented MSP species, with no oral-oriented MSP species detected for both donors, and raw saliva was dominated exclusively by oral-oriented MSP species (**Fig. 4, Fig. S5**).

**FIG 4:**
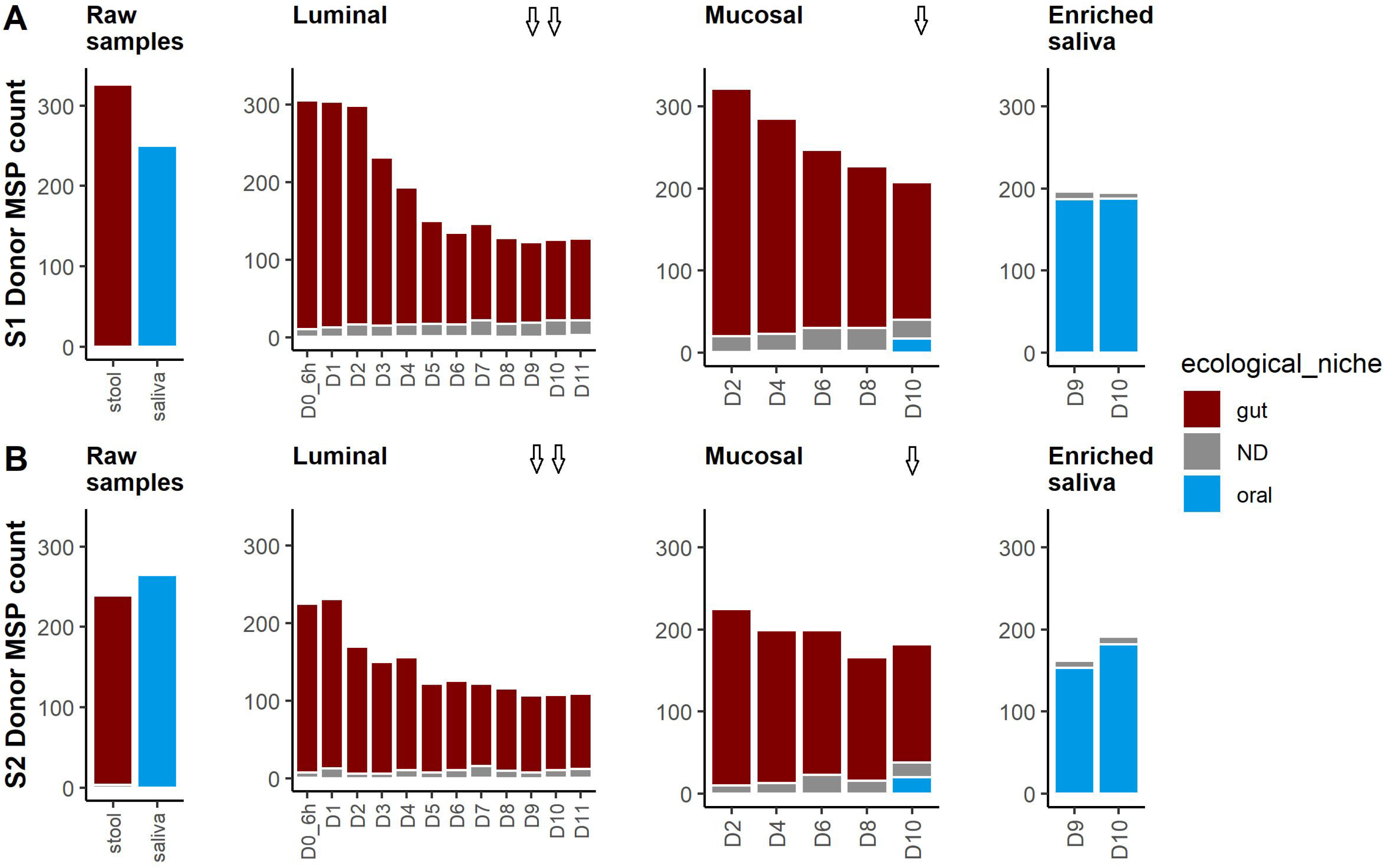
MSP species richness of oral-to-gut invasion using species ecological niche. MSP species richness was split by their corresponding ecological niche namely their dominant ecological ecosystem (brown for gut; grey for Not Determined (ND); blue for oral). **A**. S1 donor, **B.** S2 donor. Times of fermentation in the colonic M-ARCOL model are indicated in days (D). Arrows indicate saliva injection into the bioreactors on D9 and D10 days.

Following stool inoculation of the bioreactor, luminal and mucosal compartments were largely dominated by gut MSP species, yet two oral MSP species were detected even before simulated oral-to-gut invasion (**Fig. 4**). For S1 donor, these oral MSP species were detected in luminal and mucosal samples (msp_0616 *Prevotella buccae* and msp_1193 *Dialister pneumosintes*), raising to up to a fourth of the relative abundance at D8 in the luminal sample but remaining at low relative abundance (0.01) in the mucosal sample before oral-to-gut invasion. For S2 donor, only one msp_0616 *Prevotella buccae* species was detected in the luminal compartment. (**Fig. S5**). Oral MSP species were non-dominant, either by number or relative abundance, at D8 in the mucosal compartment of the 2 donors before oral-to-gut invasion.

### Preference of oral microbial invaders for the mucosal microenvironment

After the injection of enriched saliva in the system, we found only three oral MSP species in the luminal compartments of both donors at days D9 and D10 (msp_0616 *Prevotella buccae*, msp_0677c *Slackia exigua* and msp_0884 *Veillonella atypica*) (**Fig. 5**); these low numbers persisted at D11. In contrast, a much higher number, n=28, was detected in the mucosal microenvironment of both donors, representing about 15% of the enriched saliva microbial diversity. Similar numbers of oral MSP species were found for S1 and S2 donors in the mucosal samples (17 and 20 oral MSP species, respectively), but their relative abundance differed between donors (relative abundance of 0.13 and 0.0043, respectively, **Fig. S5**). These oral species invaders belonged to a limited number of taxa, with members of the *Veillonella*, *Streptococcus, Prevotella* and *Haemophilus* genera, all common taxa of the oral microbiome (**Fig. 5**). Additional common taxa included *Porphymoromas*, *Neisseria* and *Rothia* genera. We also observed that the relative abundance of these 28 oral invaders in the mucosal microenvironment was not systematically related to their respective abundance in enriched saliva samples (rho spearman 0.21 and 0.88 for S1 and S2 donors respectively, between enriched saliva and mucosal at D10; p-values = 0.28 and < 1.5e-9).

**FIG 5:**
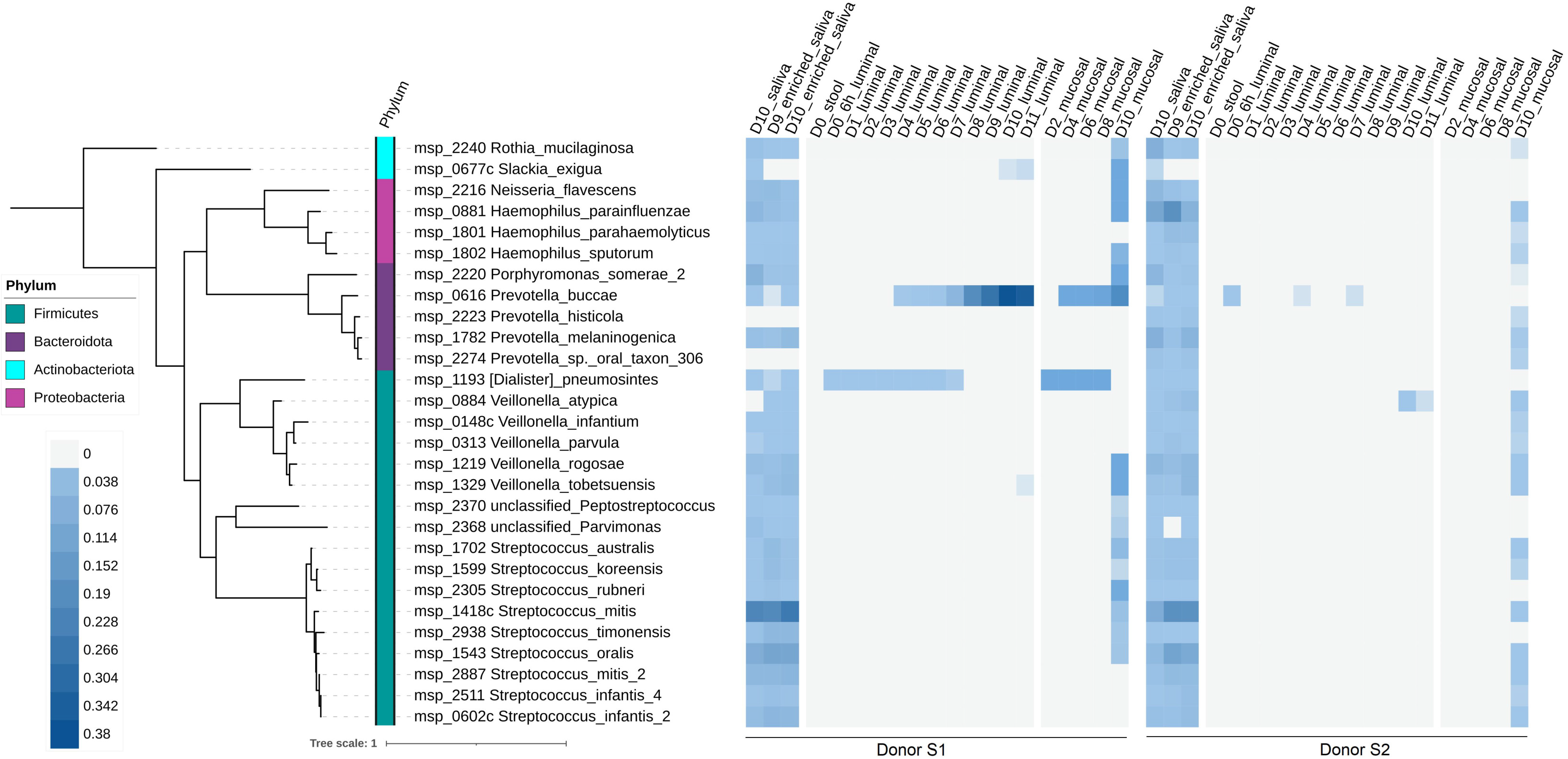
Phylogenetic tree for oral MSP species during oral-to-gut invasion. Phylogenetic tree for 28 oral MSP species detected in the initial stools, the luminal and mucosal samples from M-ARCOL, regardless of the raw and enriched saliva composition. The tree was generated using the 40 universal marker proteins from the oral MSP species extracted by the fetchMG software. For each S1 and S2 donors, a heatmap for the oral MSP species normalized abundance is reported, with a colour shade from grey (not detected) to dark blue (highly detected, maximum relative abundance = 0.38). Times of fermentation in the colonic M-ARCOL model are indicated in days (D).

## DISCUSSION

In this study, we present a new model for oral to gut microbial invasion simulation using the M-ARCOL bioreactor and whole metagenome sequencing, with the goal to provide a tool for a comprehensive understanding of the interactions of these two microbial ecosystems of the gastro-intestinal tract. This experimental set-up was motivated by the growing body of evidence for the oral microbial communities’ impact on the gut microbiome composition and its potential effect on human health (22, 24, 33, 34). The new model consists of a colonic bioreactor inoculated with faecal samples from healthy adult donors, used to follow daily microbial changes for 11 days, combined with a simulated oral invasion by injecting enriched saliva on days D9 to D10. A shotgun metagenomic based analysis was essential to achieve species level resolution, to differentiate closely related species originating from the oral or gut ecosystems. This study also required a combined salivary and faecal *in silico* microbial exploration of two healthy donors with distinct faecal microbial composition; Firmicutes dominance for S1 donor and Bacteroidetes dominance for S2 donor. Since faecal microbiota inter-individual variability is important, this pilot study was carried out on faecal samples collected from two healthy donors known to represent different conditions in terms of faecal microbial composition, methane-producing status, ages (S1: 22 years, S2: 52 years) and dietary habits (S1: neo-vegetarian diet, S2: omnivorous diet). Lastly, we assessed the oral to gut microbial interactions using the mucosal configuration of the ARCOL *in vitro* gut model (28), providing a unique opportunity to independently investigate luminal and mucosal-associated microbial communities, thanks to a distinct capture of the fine-scale regionalization of the human colon (28).

One key aspect of our investigation was to retrieve enough salivary material to perform both the oral injection into the bioreactor and the microbial DNA extraction for shotgun metagenomic analysis. To circumvent a relatively low amount of saliva biomass, we performed a microbial saliva enrichment step prior to oral injection. While this enrichment reduced somewhat the oral microbial richness, a major part was retained (70%), with a composition close to the raw saliva, thus preserving the oral microbial signatures of the two donors. Interestingly, we found that, whatever the donor, the mucosal compartment of M-ARCOL system enabled the subsistence of a higher microbial richness compared to the luminal one (28, 35, 36). This highlights the critical contribution of the mucosal set-up in microbial dynamic analysis based on colonic *in vitro* models and supports the underestimated role of the mucus in many physiological and pathological processes involving gut microbiome (37–39).

Using the microbial species ecological niche predisposition (gut or oral), we explored the number and amount of oral microbial species in the luminal or mucosal compartments of M-ARCOL before and after simulated oral invasion. As expected, variations of oral microbial species after invasion were detected, indicating that the protocol design was successful in confronting the two microbial ecosystems. While the two donors displayed different raw oral and faecal microbial compositions, it was observed that oral invasion mostly occurred in the mucosal compartment and was limited in the luminal one for both donors. This suggests invasive oral species preference for the intestinal mucosal resources. Interestingly, the oral invaders were all members of common salivary microbiota, as part of the healthy oral core microbiome (40–43). These results suggest that some oral species might possess functions to utilize MUC-2 proteins from the gut (44). We also observed for the S1 donor a significant increase and resilience of several oral species prior to the simulated oral invasion, indicating that sub-abundant oral species present in stool samples could surge when appropriate conditions occurred, such as gut microbial richness decrease (19, 45).

The main objectives of the current study were to test the feasibility of the experimental approach to simulate oral microbial invasion in the intestinal compartment and the efficiency of the saliva enrichment. Our study demonstrates promising opportunities to (i) further understand mechanisms of oral microbial invasion of the human gut microbiome (5, 46), (ii) define microbe-microbe interactions in a compartmentalized fashion (luminal *versus* mucosal) and (iii) eventually help to clarify the potential impact of oral invasion on human health (20, 47). Future developments could include the implementation of an upper *in vitro* human digestive tract by coupling the M-ARCOL with the TIM-1 stomach and small intestinal digester (30, 48, 49). This would allow the combined determination of oral microbiota survival and oral-to-intestinal microbial interactions. Several studies have shown that orally derived bacteria can colonize and persist better within the gut under diseased conditions (5, 23, 24, 33). Colon *in vitro* gut models, including the M-ARCOL, can be adapted to mimic pathological situations associated to gut microbial dysbiosis, such as obesity (50, 51), irritable bowel syndrome (52) or Inflammatory Bowel Disease (53, 54). This can be performed by inoculating them with faecal samples from patients but also adapting all the nutritional and physicochemical parameters to the diseased situations (28, 49).

## MATERIAL AND METHODS

### Samples collection for bioreactor inoculation and oral invasion

Donors were selected according to their sex (one female *versus* one male), their age (23 *versus* 52 years old), their dietary habits (one eating a flexitarian based diet *versus* one consuming a western like diet) and their methane status (one producer *versus* one non-producer). The selected donors had no history of antibiotic treatment, drug or probiotic consumption for 3 months prior to sample collection. This study is a non-interventional study with no additions to usual clinical cares. Faecal samples were prepared anaerobically as previously described (28), within 6 hours after defecation. Saliva was collected from the same donor, each donor abstained from food/drink intake for 2 hours prior to sample collection. Saliva was collected twice a day (morning and afternoon) by passive drool collection method (no spitting, no blowing). Briefly, after allowing saliva to pool at the bottom of the mouth, the head was tilted forward enabling a passive flow collected (10 mL) in a sterile container.

### Fermentation in M-ARCOL

The Mucosal ARtificial COLon (M-ARCOL) is a one-stage fermentation system, used under semi-continuous conditions which simulates the physico-chemical conditions encountered in the human colon as well as the lumen- and mucus-associated human intestinal microbial ecosystem (Applikon, Schiedam, The Netherlands) (28). It consists of pH-controlled, stirred, airtight glass vessels kept under anaerobic conditions maintained by the sole activity of resident microbiota, one vessel (300 mL) mimicking the luminal associated microbiota and a second vessel containing mucin-alginate beads (total area of beads 556 cm^2^) to mimic the mucus-associated microbiota (28). The vessels were operated with an initial sparging with O_2_-free N_2_ gas and then inoculated with faecal material from donors S1 and S2, respectively. This dynamic *in vitro* model was set-up to reproduce conditions of a healthy human adult colon with a fixed temperature of 37°C, a controlled constant pH of 6.3, a stirring speed at 400 rpm, a mean retention time of 24 h, and a redox potential (Eh) of −200 mV. A sterile nutritive medium containing various sources of carbohydrates, proteins, lipids, minerals and vitamin sources was sequentially introduced into the bioreactor as described in (28, 55).

### Gas and SCFA analysis

Analysis of O_2_, CO_2_, CH_4_ and H_2_ produced during the fermentation process in the atmospheric phase of the bioreactors was performed daily to ascertain anaerobic conditions and gas composition were verified (**Fig. S1**). Gas composition was analysed using an HP 6890 gas chromatograph (Agilent Technologies, Santa Clara, USA) coupled with a TCD detector (Agilent Technologies, Santa Clara, USA). The three majors SCFAs (acetate, butyrate and propionate) were quantified in colonic samples from luminal phase by high-performance liquid chromatography (HPLC) (Elite LaChrom, Merck Hitachi, USA) coupled with a DAD diode as described in (28).

### Mucin beads and mucin compartment

Type II mucin from porcine stomach (Sigma-Aldrich, Saint-Louis, Missouri, USA) and sodium alginate (Sigma-Aldrich, Saint-Louis, Missouri, USA) were diluted in sterile distilled water, at a concentration of 5% and 2%, respectively. To produce mucin beads, the mucin-alginate solution was dropped using a peristaltic pump into a 0.2M solution of sterile CaCl_2_ under agitation. Mucin-alginate beads were introduced in airtight glass compartment allowing the circulation of the fermentation medium and ensuring the contact of the resident luminal microbiota with the mucin beads. Mucin beads were kept at 37°C through the experiment using a hot water bath and replaced every 2 days by fresh sterile ones under a constant flow of CO_2_ to retain anaerobiosis.

### Experimental design and sampling

Following faecal inoculation of the bioreactor, fermentation was conducted for a total duration of 11 days. On days D9 and D10, 10ml of enriched saliva was introduced into the bioreactor twice a day (morning and late afternoon) after an enrichment of 9 hours. A saliva washout was realized on day D11 without the injection of enriched saliva samples. For saliva enrichment, 10mL of freshly collected saliva was resuspended in 15 mL of colonic nutritive medium (37°C, anaerobically, 100 rpm) (28) to favour oral bacteria multiplication and enrich the microbial fraction over the human fraction in the saliva samples. Circulating medium from the bioreactor vessel, referred to as luminal microbiota, was collected daily. Medium circulating in the mucin beads compartment, referred to as mucosal microbiota was collected every 2 days. The remaining late afternoon enriched saliva samples (~5 mL) were collected for each donor (referred to as enriched saliva). On day D10, a saliva sampling was performed per donor without enrichment (referred to as raw saliva).

### DNA extraction procedures

Prior to extraction, all samples were handled in the laboratory following the IHMS SOP 004 (http://www.human-microbiome.org/), and stored at −80°C. Faecal and luminal samples were aliquoted into 200 mg and DNA extraction was performed following the IHMS SOP P7 V2. A specific DNA preparation to remove human related DNA was performed on saliva and enriched saliva as follow: 200 μL of saliva was centrifuged 10 min at 10 000g for 4min before adding 190 mL of sterile Milli-Q water to the pellet. After 5 min of incubation at room temperature, 10 μL of Propidium Monoazide (Clinisciences, Nanterre, France) was added to the tube to a final concentration of 10 μM. After 5 min of incubation at room temperature, samples were kept on ice and were exposed to halogen light for 25 min. Samples were stored at −80°C until extraction following the IHMS SOP P7 V2. For mucosal samples, the same preparatory procedure was performed followed by DNA extraction using the IHMS SOP 06 V2 protocol. DNA was quantified using Qubit Fluorometric Quantitation (ThermoFisher Scientific, Waltham, USA) and qualified using DNA size profiling on a Fragment Analyzer (Agilent Technologies, Santa Clara, USA).

### High throughput sequencing

For each sample, 1 μg of high molecular weight DNA (>10 kbp) was used to build the library. DNA was shared into 150 bp fragments using an ultra-sonicator (Covaris, Woburn, US) and DNA fragment library construction was performed using the Ion Plus Fragment Library and Ion Xpress Barcode Adaptaters Kits (ThermoFisher Scientific, Waltham, USA). We used Ion Proton Sequencer and Ion GeneStudio S5 Prime Sequencer to sequence the librairies (ThermoFisher Scientific, Waltham, USA), with a minimum of 20 million 150 bp high-quality reads generated per library for luminal, mucosal and enriched saliva samples.

### Read Mapping

An average of 23.7 ± 4.5 million reads was produced. We performed quality filtering using AlienTrimmer software to discard low quality reads. Remaining human-related reads (0.02% for stools, luminal and mucosal samples and 61% for saliva and enriched-saliva, on average) were removed using Bowtie2 (56), with at least 90% identity with Human genome reference GRCh38 (accession GCA_000001405.15). Resulting high-quality reads were mapped onto the 10.4 million gut IGC2 (Integrated Gene Catalogue) catalog of the human microbiome (57) and onto the 8.4 million oral catalog (58) using the METEOR software (https://forgemia.inra.fr/metagenopolis/meteor/-/tree/master/meteor-pipeline). Mapping was performed using an identity threshold of 90% to the reference gene catalog with Bowtie2 in a two-step procedure using a downsizing level of 12 million reads per sample, as described in Meslier *et al*. (59).

### Metagenomic Species Pangenome (MSP species) determination and ecological niche definition

Metagenomic Species Pangenome (MSP) were used to identify and quantify microbial species associated to the 10.4 million human gut genes on one hand (doi:10.15454/FLANUP) and the 8.4 million human oral genes catalogs on the other hand (doi: 10.15454/WQ4UTV). MSP species are clusters of co-abundant genes (minimum cluster size > 100 genes) used as proxy for microbial species (60, 61). MSP abundances were estimated as the mean abundance of their 100 marker genes in both gut and oral catalogs, as far as at least 10% of these genes were detected (abundance strictly positive). From the independent gut and oral abundance tables, we computed a single abundance table by filtering overlapping MSP species between the two tables. MSP species ecological niche was determined by evaluating MSP detection in stools and raw saliva samples from the two donors. We assigned each MSP to gut and oral ecological niches when strict occurrence was found in either gut or saliva and not-determined (ND) when the MSP was detected in raw saliva and stool sample in at least on sample (**Table S2**). MSP species richness was determined by counting the number of MSP species detected in the corresponding sample on the merged abundance table.

### Computational analysis

All further steps were performed using R 3.5.0 (https://www.r-project.org). Data were processed and visualized using R packages *dplyr, stringr, tidyverse, ggpubr* and *pheatmap*.

## Data availability

All sequencing data have been deposited at the European Nucleotide Archive database under the study accession PRJEB52431. Associated metadata are provided in the Table S1.

## ACKNOWLEDGEMENTS

This work was supported by the European FP7 Marie Skłodowska-Curie actions AgreenSkillsPlus PCOFUND-GA-2013-609398 grant to MA. Additional funding was from the Metagenopolis grant ANR-11-DPBS-0001 and from the SCUSI OBFIBRE program from Auvergne Rhône Alpes Region to MEDIS.

## Contributions

Oral invasion experiment design: MA, SDE; M-ARCOL experiment design: SBD, LEM; Bioreactor monitoring and sampling: LEM, OU, CD, EF and SD; DNA extraction: AD, SJ, CM; Metagenomic sequencing: BQ; Metagenomic data pre-processing: VM, FT; Oral microbial catalog providers: FPO and ELC; Data analysis and Figures: VM and MA; Data interpretation: VM, MA, LEM, SBD; Writing VM, LEM, MA and SBD; Review and editing: VM, LEM, MA and SBD; Resources: MA and SBD.

## Competing Interests

The authors declare no conflict of interests.

## Ethics declarations

This study is a non-interventional study with no additions to usual clinical cares. The donors provided a written consent for the analysis and publication of their oral and fecal samples in the specific context of this study. This non-interventional study did not require approval from an ethics committee according to the French Public Health Law (Code de la santé publique article L 1121-1.1).

## REFERENCES

1. Qin J, Li R, Raes J, Arumugam M, Burgdorf KS, Manichanh C, Nielsen T, Pons N, Levenez F, Yamada T, Mende DR, Li J, Xu J, Li S, Li D, Cao J, Wang B, Liang H, Zheng H, Xie Y, Tap J, Lepage P, Bertalan M, Batto J-M, Hansen T, Le Paslier D, Linneberg A, Nielsen HB, Pelletier E, Renault P, Sicheritz-Ponten T, Turner K, Zhu H, Yu C, Li S, Jian M, Zhou Y, Li Y, Zhang X, Li S, Qin N, Yang H, Wang J, Brunak S, Doré J, Guarner F, Kristiansen K, Pedersen O, Parkhill J, Weissenbach J, MetaHIT Consortium, Bork P, Ehrlich SD, Wang J. 2010. A human gut microbial gene catalogue established by metagenomic sequencing. Nature 464:59–65.

2. Fan Y, Pedersen O. 2021. Gut microbiota in human metabolic health and disease. Nat Rev Microbiol 19:55–71.

3. Lynch SV, Pedersen O. 2016. The Human Intestinal Microbiome in Health and Disease. N Engl J Med 375:2369–2379.

4. NIH HMP Working Group, Peterson J, Garges S, Giovanni M, McInnes P, Wang L, Schloss JA, Bonazzi V, McEwen JE, Wetterstrand KA, Deal C, Baker CC, Di Francesco V, Howcroft TK, Karp RW, Lunsford RD, Wellington CR, Belachew T, Wright M, Giblin C, David H, Mills M, Salomon R, Mullins C, Akolkar B, Begg L, Davis C, Grandison L, Humble M, Khalsa J, Little AR, Peavy H, Pontzer C, Portnoy M, Sayre MH, Starke-Reed P, Zakhari S, Read J, Watson B, Guyer M. 2009. The NIH Human Microbiome Project. Genome Res 19:2317–2323.

5. Park S-Y, Hwang B-O, Lim M, Ok S-H, Lee S-K, Chun K-S, Park K-K, Hu Y, Chung W-Y, Song N-Y. 2021. Oral-Gut Microbiome Axis in Gastrointestinal Disease and Cancer. Cancers (Basel) 13:2124.

6. Human Microbiome Project Consortium. 2012. Structure, function and diversity of the healthy human microbiome. Nature 486:207–214.

7. Dawes C. 2008. Salivary flow patterns and the health of hard and soft oral tissues. J Am Dent Assoc 139 Suppl:18S–24S.

8. Sedghi L, DiMassa V, Harrington A, Lynch SV, Kapila YL. 2021. The oral microbiome: Role of key organisms and complex networks in oral health and disease. Periodontol 2000 87:107–131.

9. Dewhirst FE, Chen T, Izard J, Paster BJ, Tanner ACR, Yu W-H, Lakshmanan A, Wade WG. 2010. The human oral microbiome. J Bacteriol 192:5002–5017.

10. Zhao H, Chu M, Huang Z, Yang X, Ran S, Hu B, Zhang C, Liang J. 2017. Variations in oral microbiota associated with oral cancer. Sci Rep 7:11773.

11. Segata N, Haake SK, Mannon P, Lemon KP, Waldron L, Gevers D, Huttenhower C, Izard J. 2012. Composition of the adult digestive tract bacterial microbiome based on seven mouth surfaces, tonsils, throat and stool samples. Genome Biol 13:R42.

12. Roberts JA. 1996. Tropism in Bacterial Infections: Urinary Tract Infections. Journal of Urology 156:1552–1559.

13. Pereira FC, Berry D. 2017. Microbial nutrient niches in the gut. Environ Microbiol 19:1366–1378.

14. Vuik F, Dicksved J, Lam SY, Fuhler GM, van der Laan L, van de Winkel A, Konstantinov SR, Spaander M, Peppelenbosch MP, Engstrand L, Kuipers EJ. 2019. Composition of the mucosa-associated microbiota along the entire gastrointestinal tract of human individuals. United European Gastroenterol J 7:897–907.

15. Wu GD, Chen J, Hoffmann C, Bittinger K, Chen Y-Y, Keilbaugh SA, Bewtra M, Knights D, Walters WA, Knight R, Sinha R, Gilroy E, Gupta K, Baldassano R, Nessel L, Li H, Bushman FD, Lewis JD. 2011. Linking long-term dietary patterns with gut microbial enterotypes. Science 334:105–108.

16. Arumugam M, Raes J, Pelletier E, Le Paslier D, Yamada T, Mende DR, Fernandes GR, Tap J, Bruls T, Batto J-M, Bertalan M, Borruel N, Casellas F, Fernandez L, Gautier L, Hansen T, Hattori M, Hayashi T, Kleerebezem M, Kurokawa K, Leclerc M, Levenez F, Manichanh C, Nielsen HB, Nielsen T, Pons N, Poulain J, Qin J, Sicheritz-Ponten T, Tims S, Torrents D, Ugarte E, Zoetendal EG, Wang J, Guarner F, Pedersen O, de Vos WM, Brunak S, Doré J, MetaHIT Consortium, Antolín M, Artiguenave F, Blottiere HM, Almeida M, Brechot C, Cara C, Chervaux C, Cultrone A, Delorme C, Denariaz G, Dervyn R, Foerstner KU, Friss C, van de Guchte M, Guedon E, Haimet F, Huber W, van Hylckama-Vlieg J, Jamet A, Juste C, Kaci G, Knol J, Lakhdari O, Layec S, Le Roux K, Maguin E, Mérieux A, Melo Minardi R, M’rini C, Muller J, Oozeer R, Parkhill J, Renault P, Rescigno M, Sanchez N, Sunagawa S, Torrejon A, Turner K, Vandemeulebrouck G, Varela E, Winogradsky Y, Zeller G, Weissenbach J, Ehrlich SD, Bork P. 2011. Enterotypes of the human gut microbiome. Nature 473:174–180.

17. Almeida A, Mitchell AL, Boland M, Forster SC, Gloor GB, Tarkowska A, Lawley TD, Finn RD. 2019. A new genomic blueprint of the human gut microbiota. Nature 568:499–504.

18. Lloyd-Price J, Abu-Ali G, Huttenhower C. 2016. The healthy human microbiome. Genome Med 8:51.

19. Le Chatelier E, Nielsen T, Qin J, Prifti E, Hildebrand F, Falony G, Almeida M, Arumugam M, Batto J-M, Kennedy S, Leonard P, Li J, Burgdorf K, Grarup N, Jørgensen T, Brandslund I, Nielsen HB, Juncker AS, Bertalan M, Levenez F, Pons N, Rasmussen S, Sunagawa S, Tap J, Tims S, Zoetendal EG, Brunak S, Clément K, Doré J, Kleerebezem M, Kristiansen K, Renault P, Sicheritz-Ponten T, de Vos WM, Zucker J-D, Raes J, Hansen T, MetaHIT consortium, Bork P, Wang J, Ehrlich SD, Pedersen O. 2013. Richness of human gut microbiome correlates with metabolic markers. Nature 500:541–546.

20. Li B, Ge Y, Cheng L, Zeng B, Yu J, Peng X, Zhao J, Li W, Ren B, Li M, Wei H, Zhou X. 2019. Oral bacteria colonize and compete with gut microbiota in gnotobiotic mice. Int J Oral Sci 11:10.

21. Schmidt TS, Hayward MR, Coelho LP, Li SS, Costea PI, Voigt AY, Wirbel J, Maistrenko OM, Alves RJ, Bergsten E, de Beaufort C, Sobhani I, Heintz-Buschart A, Sunagawa S, Zeller G, Wilmes P, Bork P. 2019. Extensive transmission of microbes along the gastrointestinal tract. Elife 8:e42693.

22. Hu S, Png E, Gowans M, Ong DEH, de Sessions PF, Song J, Nagarajan N. 2021. Ectopic gut colonization: a metagenomic study of the oral and gut microbiome in Crohn’s disease. Gut Pathog 13:13.

23. Read E, Curtis MA, Neves JF. 2021. The role of oral bacteria in inflammatory bowel disease. Nat Rev Gastroenterol Hepatol 18:731–742.

24. Qin N, Yang F, Li A, Prifti E, Chen Y, Shao L, Guo J, Le Chatelier E, Yao J, Wu L, Zhou J, Ni S, Liu L, Pons N, Batto JM, Kennedy SP, Leonard P, Yuan C, Ding W, Chen Y, Hu X, Zheng B, Qian G, Xu W, Ehrlich SD, Zheng S, Li L. 2014. Alterations of the human gut microbiome in liver cirrhosis. Nature 513:59–64.

25. Farin W, Oñate FP, Plassais J, Bonny C, Beglinger C, Woelnerhanssen B, Nocca D, Magoules F, Le Chatelier E, Pons N, Cervino ACL, Ehrlich SD. 2020. Impact of laparoscopic Roux-en-Y gastric bypass and sleeve gastrectomy on gut microbiota: a metagenomic comparative analysis. Surg Obes Relat Dis 16:852–862.

26. Van den Abbeele P, Grootaert C, Marzorati M, Possemiers S, Verstraete W, Gérard P, Rabot S, Bruneau A, El Aidy S, Derrien M, Zoetendal E, Kleerebezem M, Smidt H, Van de Wiele T. 2010. Microbial community development in a dynamic gut model is reproducible, colon region specific, and selective for Bacteroidetes and Clostridium cluster IX. Appl Environ Microbiol 76:5237–5246.

27. Zihler Berner A, Fuentes S, Dostal A, Payne AN, Vazquez Gutierrez P, Chassard C, Grattepanche F, de Vos WM, Lacroix C. 2013. Novel Polyfermentor intestinal model (PolyFermS) for controlled ecological studies: validation and effect of pH. PLoS One 8:e77772.

28. Deschamps C, Fournier E, Uriot O, Lajoie F, Verdier C, Comtet-Marre S, Thomas M, Kapel N, Cherbuy C, Alric M, Almeida M, Etienne-Mesmin L, Blanquet-Diot S. 2020. Comparative methods for fecal sample storage to preserve gut microbial structure and function in an in vitro model of the human colon. Appl Microbiol Biotechnol 104:10233–10247.

29. Daniel N, Lécuyer E, Chassaing B. 2021. Host/microbiota interactions in health and diseases-Time for mucosal microbiology!Mucosal Immunol 14:1006–1016.

30. Etienne-Mesmin L, Chassaing B, Desvaux M, De Paepe K, Gresse R, Sauvaitre T, Forano E, de Wiele TV, Schüller S, Juge N, Blanquet-Diot S. 2019. Experimental models to study intestinal microbes-mucus interactions in health and disease. FEMS Microbiol Rev 43:457–489.

31. Sauvaitre T, Etienne-Mesmin L, Sivignon A, Mosoni P, Courtin CM, Van de Wiele T, Blanquet-Diot S. 2021. Tripartite relationship between gut microbiota, intestinal mucus and dietary fibers: towards preventive strategies against enteric infections. FEMS Microbiol Rev 45:fuaa052.

32. Van den Abbeele P, Roos S, Eeckhaut V, MacKenzie DA, Derde M, Verstraete W, Marzorati M, Possemiers S, Vanhoecke B, Van Immerseel F, Van de Wiele T. 2012. Incorporating a mucosal environment in a dynamic gut model results in a more representative colonization by lactobacilli. Microb Biotechnol 5:106–115.

33. Flemer B, Warren RD, Barrett MP, Cisek K, Das A, Jeffery IB, Hurley E, O’Riordain M, Shanahan F, O’Toole PW. 2018. The oral microbiota in colorectal cancer is distinctive and predictive. Gut 67:1454–1463.

34. Fan X, Alekseyenko AV, Wu J, Peters BA, Jacobs EJ, Gapstur SM, Purdue MP, Abnet CC, Stolzenberg-Solomon R, Miller G, Ravel J, Hayes RB, Ahn J. 2018. Human oral microbiome and prospective risk for pancreatic cancer: a population-based nested case-control study. Gut 67:120–127.

35. Duncan K, Carey-Ewend K, Vaishnava S. 2021. Spatial analysis of gut microbiome reveals a distinct ecological niche associated with the mucus layer. Gut Microbes 13:1874815.

36. Li H, Limenitakis JP, Fuhrer T, Geuking MB, Lawson MA, Wyss M, Brugiroux S, Keller I, Macpherson JA, Rupp S, Stolp B, Stein JV, Stecher B, Sauer U, McCoy KD, Macpherson AJ. 2015. The outer mucus layer hosts a distinct intestinal microbial niche. Nat Commun 6:8292.

37. Corfield AP. 2018. The Interaction of the Gut Microbiota with the Mucus Barrier in Health and Disease in Human. Microorganisms 6:E78.

38. Grondin JA, Kwon YH, Far PM, Haq S, Khan WI. 2020. Mucins in Intestinal Mucosal Defense and Inflammation: Learning From Clinical and Experimental Studies. Front Immunol 11:2054.

39. Paone P, Cani PD. 2020. Mucus barrier, mucins and gut microbiota: the expected slimy partners? Gut 69:2232–2243.

40. Zaura E, Keijser BJF, Huse SM, Crielaard W. 2009. Defining the healthy “core microbiome” of oral microbial communities. BMC Microbiol 9:259.

41. Liu G, Tang CM, Exley RM. 2015. Non-pathogenic Neisseria: members of an abundant, multi-habitat, diverse genus. Microbiology (Reading) 161:1297–1312.

42. van der Hoeven JS, van den Kieboom CW, Camp PJ. 1990. Utilization of mucin by oral Streptococcus species. Antonie Van Leeuwenhoek 57:165–172.

43. Derrien M, van Passel MW, van de Bovenkamp JH, Schipper RG, de Vos WM, Dekker J. 2010. Mucin-bacterial interactions in the human oral cavity and digestive tract. Gut Microbes 1:254–268.

44. Kasprzak A, Adamek A. 2019. Mucins: the Old, the New and the Promising Factors in Hepatobiliary Carcinogenesis. Int J Mol Sci 20:E1288.

45. Solé C, Guilly S, Da Silva K, Llopis M, Le-Chatelier E, Huelin P, Carol M, Moreira R, Fabrellas N, De Prada G, Napoleone L, Graupera I, Pose E, Juanola A, Borruel N, Berland M, Toapanta D, Casellas F, Guarner F, Doré J, Solà E, Ehrlich SD, Ginès P. 2021. Alterations in Gut Microbiome in Cirrhosis as Assessed by Quantitative Metagenomics: Relationship With Acute-on-Chronic Liver Failure and Prognosis. Gastroenterology 160:206–218.e13.

46. Kitamoto S, Nagao-Kitamoto H, Jiao Y, Gillilland MG, Hayashi A, Imai J, Sugihara K, Miyoshi M, Brazil JC, Kuffa P, Hill BD, Rizvi SM, Wen F, Bishu S, Inohara N, Eaton KA, Nusrat A, Lei YL, Giannobile WV, Kamada N. 2020. The Intermucosal Connection between the Mouth and Gut in Commensal Pathobiont-Driven Colitis. Cell 182:447–462.e14.

47. Khor B, Snow M, Herrman E, Ray N, Mansukhani K, Patel KA, Said-Al-Naief N, Maier T, Machida CA. 2021. Interconnections Between the Oral and Gut Microbiomes: Reversal of Microbial Dysbiosis and the Balance Between Systemic Health and Disease. Microorganisms 9:496.

48. Roussel C, De Paepe K, Galia W, de Bodt J, Chalancon S, Denis S, Leriche F, Vandekerkove P, Ballet N, Blanquet-Diot S, Van de Wiele T. Multi-targeted properties of the probiotic saccharomyces cerevisiae CNCM I-3856 against enterotoxigenic escherichia coli (ETEC) H10407 pathogenesis across human gut models. Gut Microbes 13:1953246.

49. Fournier E, Etienne-Mesmin L, Grootaert C, Jelsbak L, Syberg K, Blanquet-Diot S, Mercier-Bonin M. 2021. Microplastics in the human digestive environment: A focus on the potential and challenges facing in vitro gut model development. J Hazard Mater 415:125632.

50. Uriot O, Deschamps C, Brun M, Pouget M, Alric M, Chaudemanche C, Boirie Y, Etienne-Mesmin L, Blanquet-Diot S. 2022. Development and validation of an in vitro colonic model of gut microbiota dysbiosis associated to obesity. Cork, Ireland.

51. Aguirre M, Bussolo de Souza C, Venema K. 2016. The Gut Microbiota from Lean and Obese Subjects Contribute Differently to the Fermentation of Arabinogalactan and Inulin. PLoS One 11:e0159236.

52. Uriot O, Fournier E, Lajoie F, Kerchove N, Scanzy J, Denis S, Alric M, Etienne-Mesmin L, Blanquet-Diot S. 2021. Development of an in vitro model of dysbiotic colonic conditions of IBS-D patients. Clermont-Ferrand, France.

53. Vermeiren J, Van den Abbeele P, Laukens D, Vigsnaes LK, De Vos M, Boon N, Van de Wiele T. 2012. Decreased colonization of fecal Clostridium coccoides/Eubacterium rectale species from ulcerative colitis patients in an in vitro dynamic gut model with mucin environment. FEMS Microbiol Ecol 79:685–696.

54. Van den Abbeele P, Marzorati M, Derde M, De Weirdt R, Joan V, Possemiers S, Van de Wiele T. 2016. Arabinoxylans, inulin and Lactobacillus reuteri 1063 repress the adherent-invasive Escherichia coli from mucus in a mucosa-comprising gut model. NPJ Biofilms Microbiomes 2:16016.

55. Thévenot J, Cordonnier C, Rougeron A, Le Goff O, Nguyen HTT, Denis S, Alric M, Livrelli V, Blanquet-Diot S. 2015. Enterohemorrhagic Escherichia coli infection has donor-dependent effect on human gut microbiota and may be antagonized by probiotic yeast during interaction with Peyer’s patches. Appl Microbiol Biotechnol 99:9097–9110.

56. Langmead B, Salzberg SL. 2012. Fast gapped-read alignment with Bowtie 2. Nat Methods 9:357–359.

57. Wen C, Zheng Z, Shao T, Liu L, Xie Z, Le Chatelier E, He Z, Zhong W, Fan Y, Zhang L, Li H, Wu C, Hu C, Xu Q, Zhou J, Cai S, Wang D, Huang Y, Breban M, Qin N, Ehrlich SD. 2017. Quantitative metagenomics reveals unique gut microbiome biomarkers in ankylosing spondylitis. Genome Biol 18:142.

58. Le Chatelier E, Almeida M, Plaza Oñate F, Pons N, Gauthier F, Ghozlane A, Ehrlich SD, Witherden E, Gomez-Cabrero D. 2021. A catalog of genes and species of the human oral microbiota. Portail Data INRAE.

59. Meslier V, Laiola M, Roager HM, De Filippis F, Roume H, Quinquis B, Giacco R, Mennella I, Ferracane R, Pons N, Pasolli E, Rivellese A, Dragsted LO, Vitaglione P, Ehrlich SD, Ercolini D. 2020. Mediterranean diet intervention in overweight and obese subjects lowers plasma cholesterol and causes changes in the gut microbiome and metabolome independently of energy intake. Gut 69:1258–1268.

60. Plaza Oñate F, Le Chatelier E, Almeida M, Cervino ACL, Gauthier F, Magoulès F, Ehrlich SD, Pichaud M. 2019. MSPminer: abundance-based reconstitution of microbial pan-genomes from shotgun metagenomic data. Bioinformatics 35:1544–1552.

61. Nielsen HB, Almeida M, Juncker AS, Rasmussen S, Li J, Sunagawa S, Plichta DR, Gautier L, Pedersen AG, Le Chatelier E, Pelletier E, Bonde I, Nielsen T, Manichanh C, Arumugam M, Batto J-M, Quintanilha Dos Santos MB, Blom N, Borruel N, Burgdorf KS, Boumezbeur F, Casellas F, Doré J, Dworzynski P, Guarner F, Hansen T, Hildebrand F, Kaas RS, Kennedy S, Kristiansen K, Kultima JR, Léonard P, Levenez F, Lund O, Moumen B, Le Paslier D, Pons N, Pedersen O, Prifti E, Qin J, Raes J, Sørensen S, Tap J, Tims S, Ussery DW, Yamada T, MetaHIT Consortium, Renault P, Sicheritz-Ponten T, Bork P, Wang J, Brunak S, Ehrlich SD, MetaHIT Consortium. 2014. Identification and assembly of genomes and genetic elements in complex metagenomic samples without using reference genomes. Nat Biotechnol 32:822–828.

